# A Graph Curvature-Based Pipeline for Discovering Immune Checkpoint Response Biomarkers

**DOI:** 10.1101/2024.09.04.611306

**Authors:** James Bannon, Charles R. Cantor, Bud Mishra

## Abstract

Immune checkpoint inhibitors (ICIs), also called immune checkpoint blockers, are a promising category of targeted therapy for solid tumors. Predicting which patients will respond to ICI therapy remains an open problem under active investigation. This paper adds to this effort by developing a modular pipeline for the discovery of biomarkers from tumor RNA-sequencing data. We contextualize gene expression measurements using a protein-protein interaction (PPI) network and use a notion of graph curvature to find (pairs of) genes in the PPI that could serve as potential biomarkers. Our candidate biomarkers are evaluated using an extensive literature search and transfer learning experiments. We also provide a harmonized collection of drug-specific candidate markers found through rank aggregation that we believe merit further study.

## 1 Introduction

Immune checkpoint inhibitors (ICIs) are a relatively new class of targeted drugs that work by re-engaging the immune system’s natural anti-tumor capabilities [1]. ICI treatment can be tremendously effective and has demonstrated longer survival times, greater tumor shrinkage, and better quality of life[2]. The downside is that not every patient benefits. Estimates of the percentage of patients who respond to therapy range between 20% and 40% [3, 2]. Worse, expression levels of the targeted proteins (e.g. PD-1, PD-L1, and CTLA-4) are inconsistent predictors of ICI response. For example, in the literature there are studies showing a positive correlation between PD-L1 expression and ICI response [4, 5] in lung cancer and others find no such association [6, 7]. Bioinformatics researchers have consequently dedicated a large amount of effort towards finding predictive biomarkers across multiple “omics” modalities like tumor-immune infiltration [8, 9, 10], tumor mutational burden [11], among many others [12]. The search for consistent biomarkers of ICI response remains an active area of research.

This paper adds to that effort by presenting a novel modular computational pipeline for the discovery of ICI response biomarkers from tumor RNA sequencing data. Following the literature we will refe to whole-tumor shotgun RNA sequencing data as ‘bulk RNA-seq’ [13]. In line with prior work [14, 15, 16], our pipeline puts gene expression measurements in the context of their biological interactions via a protein-protein interaction (PPI) network. Our pipeline uses a notion of *graph curvature*, called the Ollivier Ricci Curvature (ORC) [17], to identify edges in the PPI network that are “vulnerabilities” among responders and non-responders or vice versa [18].

ORC has been used to analyze cancer data before. The pioneering works in this field [18, 19], which form the main inspiration for our present work, analyzed data from The Cancer Genome Atlas (TCGA) [20] and used the correlation values between gene expression distributions to weight the edges in a PPI network. These weights were used to find edges which had different ORC values among cancer and normal samples. Similar efforts have been conducted across patient samples from a diverse set of cancers, as well as in the analysis of drug response in cancer cell lines, with promising results [21, 22, 23, 24, 25, 26, 22, 27].

The use of ORC to investigate the determinants of ICI response is less well explored. In fact, to our knowledge only two works exist. One paper uses the *total* curvature as a prognostic biomarker in a small cohort of patients with ovarian cancer [28]. The other uses ORC to identify gene expression modules that are correlated with ICI response [29].

Our work extends and improves upon these two papers in a number of ways. First, our approach is *tissue specific* in the way that it constructs the PPI graph. Second, ours is the largest exploration of the utility of ORC in finding ICI biomarkers to date and is also the first work to look at multiple ICI drugs applied to different tissues. Further, by focusing on gene expression our measure of gene-gene interaction is more biologically plausible than that of [28] and by focusing on patient-specific weights in a manner similar to [30] we are able to evaluate curvature differences between responders and non-responders statistically. We also are the first to test the ability of curvature-derived biomarkers to generalize beyond the dataset in which they were discovered.

We evaluate the results of our pipeline with an extensive literature review and through transfer learning experiments. We find that the pipeline recovers a number of known and recently suggested biomarkers. We also find that the biomarkers found in one patient cohort generalize to other cohorts. Finally, we use rank aggregation to propose a collection of drug-specific biomarkers that we believe are worthy of further investigation.

## 2 Results

The input to the pipeline, diagrammed in Figure 1, is a *cohort*, which we take to be a collection of *n* patients with the same tumor tissue of origin and treated with the same ICI. We assume that we have bulk RNA-seq measurement vectors ***x***^(1)^*, … ,* ***x***^(^*^n^*^)^ (taken before ICI treatment) and matched labels as responders or non-responders derived from a clinical measure such as RECIST [31]. The entry 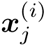 contains the expression level of gene *j* in patient *i* measured in transcripts per million (TPM). We now sketch the steps of the pipeline with details provided in Section 4.

**Figure 1:**
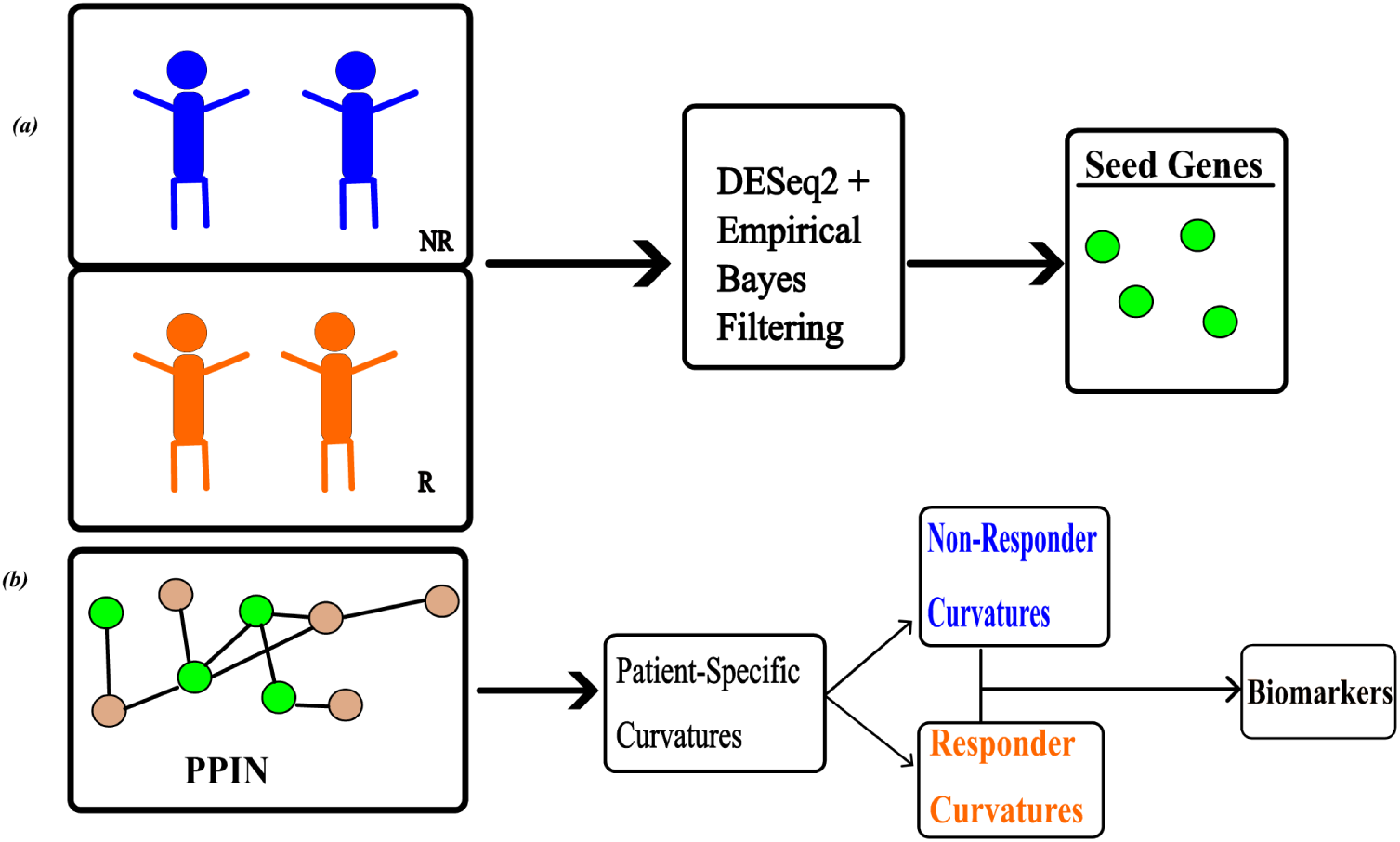
A schematic of the pipeline which is applied to each patient cohort. **(a)** For a given collection of patients we separate them into responders and non-responders. We create a collection of *seed genes* with a combination of differential expression analysis and empirical Bayes resampling. **(b)** Given the seed genes (green dots) we construct a common PPI network by looking at their neighborhoods. Lastly, for each patient we compute patient-specific edge weights and edge curvatures and select the edges with statistically significant curvature distributions between responders and non-responders.

The first step in the pipeline is the creation of a set of *seed genes*, which we determine by combining differential expression (DE) analysis and empirical Bayes (EB) techniques into a novel bootstrapping procedure. The seed genes are those genes that can be said to be differentially expressed between responders and non-responders above a stringent probability threshold. Building off of the intuition of previous studies which showed that genes close to drug targets in a PPI network have prognostic value [14, 15] we created cohort-specific PPI networks by looking around the seed genes in our network. Specifically, we chose the largest connected component of the union of the *k*-hop neighborhoods of the seed genes (see Section 4).

Then for each patient in the cohort we compute patient-specific edge weights and curvatures based on the expression vector ***x***^(^*^i^*^)^. Specifically if *r* and *s* are genes connected by an edge *e* = {*r, s*} in the cohort-specific PPI network then the weight of that edge for patient *i* is calculated as

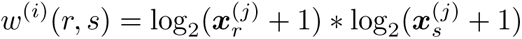

We denote the Ollivier-Ricci curvature (ORC) value of this edge for patient *i* as *κ*^(^*^i^*^)^(*r, s*) with the details of the computation given in Section 4. At the end of this process, for each edge we have a collection *K_R_*(*e*) of curvature values for the edge among responders and a similar collection *K_NR_*(*e*) among non-responders. We compared *K_R_*(*e*) to *K_NR_*(*e*) for each edge using the Mann-Whitney U-test [32] and adjusted all of the resulting *p*-values for multiple hypothesis testing with the Benjamini-Hochberg procedure [33].

For a threshold *τ* increasing in series over the values 0.005, 0.01, 0.05, and 0.10, we looked at the edges with adjusted p-values less than or equal to *τ* . The genes that were the end points of these remaining edges that were also *not* among the seed genes were considered as initial candidate biomarkers. We list these genes in Table 1 and the number of candidate biomarkers at each threshold in Appendix A. These lists were further trimmed by keeping only those genes which had different expression distributions between responders and non-responders (Mann-Whitney p-value less than 0.1 after adjustment for multiple testing). The expression distributions for each of the candidate biomarkers among responders and non-responders are displayed in Figure 2.

**Figure 2:**
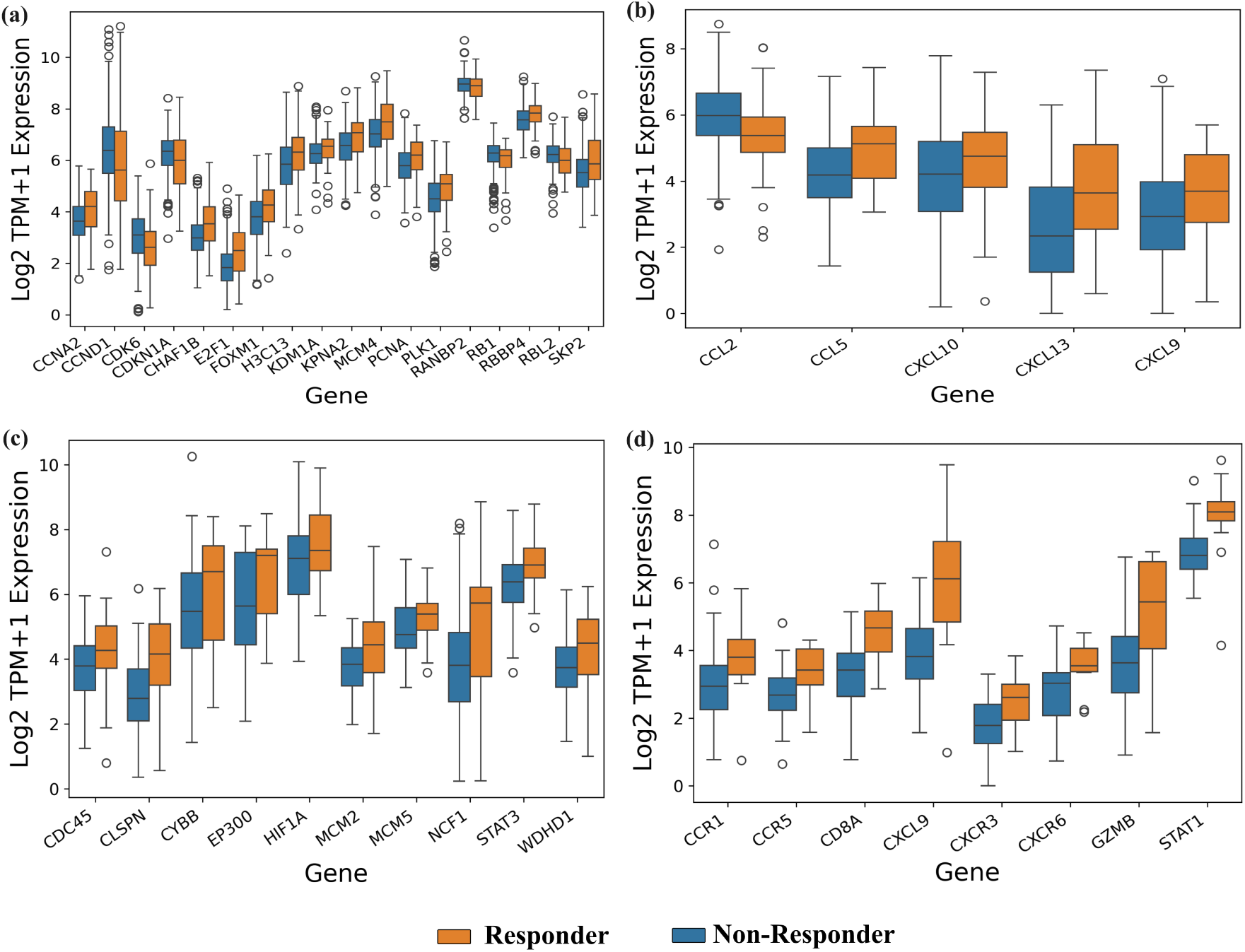
Expression distributions of candidate biomarkers in each cohort in terms of log-transformed transcripts per million (TPM) i.e: log_2_(*TPM* + 1). Candidate biomarkers consist of those genes which were not seed genes, were found to have different curvatures and had significantly different expression distributions among responders and non-responders (Mann-Whitney U p-value less than 0.1 after adjusting for multiple hypothesis testing). **(a)** Expression distributions for the candidate biomarkers for the cohort of BLCA patients treated with atezo. The genes are those derived from our pipeline with *τ* = 0.005. The remaining figures correspond to the KIRC patients treated with atezo, with *τ* = 0.01 **(b)**, the SKCM patients treated with nivo, *τ* = 0.05 **(c)** and the STAD patients treated with pembro *τ* = 0.01 **(d)**.

**Table 1:**
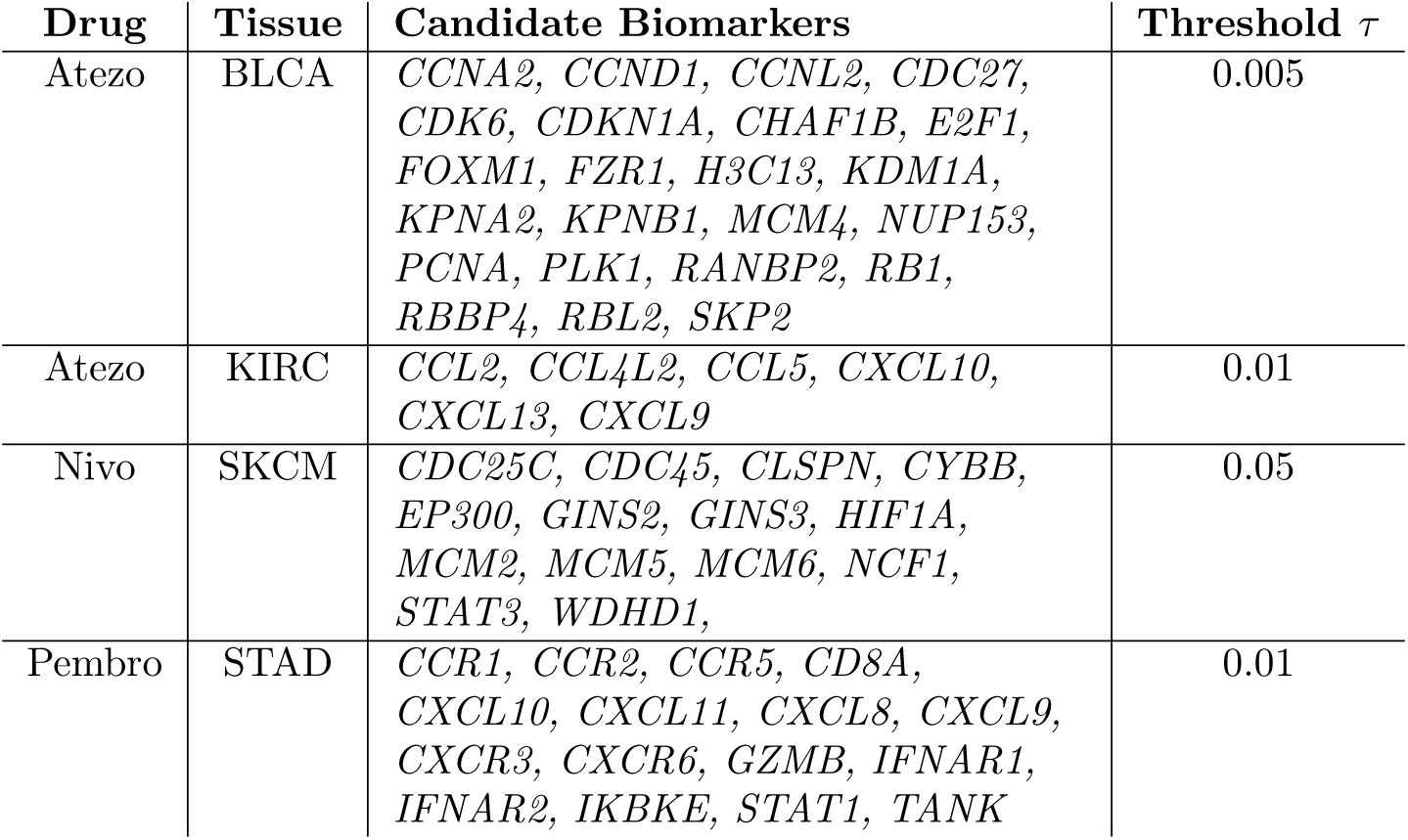
Table of the candidate biomarkers that were discovered in each cohort at the specified threshold. These gene sets were subjected to further filtering.

### 2.1 Analysis of Discovered Biomarkers

We applied our pipeline to a number of cohorts. Specifically we analyzed patients with bladder urothelial carcinoma (BLCA, n= 298) and kidney renal clear cell cancer (KIRC, 167) treated with atezolizumab (atezo), patients with skin cutaneous melanoma (SKCM, n = 109) treated with nivolumab (nivo), and patients with stomach adenocarcinoma (STAD, n=45) treated with pembrolizumab (pembro).

#### Atezo BLCA Cohort

At the threshold of *τ* = 0.005 we found 18 genes. Among these genes we found several — *CCNA2*, *CCND1*, *E2F1* — which are known to shape the tumor microenvironment. A large scale study showed that *CCNA2* was differentially expressed between cancers and normal tissue samples and played a role in how immune cells infiltrate the tumor [34]. In addition a small study suggests that *CCNA2* is a prognostic biomarker for the speed of growth in colorectal cancers [35]. Similarly *E2F1* was found to be correlated with the infiltration of multiple types of immune cells [36]. *CCND1* is known to play a role in immunosuppression and higher expression levels are associated with worse outcomes in response to ICI therapy [37], which is mirrored in the expression distributions in Figure 2.

The gene set also includes potential drug targets such as *PCNA* [38] and *CDK6* [39]. *CDK6* is of particular interest as it can enable resistance to immunotherapies [40]. *KDM1A* has also been suggested as a gene to target as part of combination therapy regimens that include ICIs [41]. *KDM1A* also upregulates the expression of atezo’s target protein PD-L1 [42].

Lastly, *FOXM1* is considered to be a so-called ’master regulator’ gene and has been documented as one of the key targetable genes in cancer [43, 44, 45].

#### Atezo KIRC Cohort

Below the threshold of 0.005 our pipeline returned four genes: *CCL5*, *CXCL10*, *CXCL13*, and *CXCL9*. Relaxing the threshold to 0.01 returned one further gene: *CCL2*. *CXCL9* is a well known ICI response biomarker which has been confirmed via a robust meta-analysis [12]. *CCL5* has also been shown to be correlated with immune cell infiltration in KIRC [46] and elevated expression of CCL5 is an ICI response biomarker in small cell lung cancer [47]. Likewise *CXCL13* is a biomarker of PD-1 therapy response in ovarian cancer due to its high correlation with immune cell infiltration [48, 12]. Finally, *CXCL10* is a known biomarker of ICI response in melanoma [49] and *CCL2* recruits immunosuppressive cells to the tumore [50].

#### Nivo SKCM Cohort

We found no edges with curvature distributions with p-values less than 0.005 or 0.01. At the threshold of 0.05 we found 10 genes, which are listed in Table 1. We only discuss a few of them here.

The most well-supported candidate biomarker genes in the collection are *STAT3* and *WDHD1*. *STAT3* is under extensive investigation as a gene to be targeted alongside ICI treatments, specifically PD-1 inhibitors [51, 52, 53, 54]. Similarly, a pan-cancer analysis suggests that *WDHD1* can be used for cancer diagnosis and prognosis in addition to predicting response to immunotherapy [55].

*CLSPN* is known to be correlated with immune cell migration into the tumor, as well as tumor mutational burden (TMB) [56], which are both known biomarkers of ICI response [11, 8]. Two other genes, *MCM2* and *NCF1* are under investigation as possible biomarkers as well [57, 58].

#### Pembro STAD Cohort

We found eight genes below the threshold of 0.01. Of these, *CCR1* has been considered as a therapeutic target for multiple myeloma [59] and, in combination with *CXCR2*, has been implicated in the way in which the tumor-immune microenvironment evolves [60]. *CCR5* is expressed by T-Cells (both CD4+ and CD8+) and boosts anti-tumor immune function [61]. Preliminary research has indicated that *CXCR3* could be used as a prognostic biomarker in part because it has other ICI biomarkers — *CXCL9* and *CXCL10* — among its ligands [62, 12]. Similarly *STAT1* expression levels have shown early promise as a biomarker for ICI therapy as its expression is correlated with the expression of PD-L1 [63, 64]. There is also literature suggesting that *CD8A* and *GZMB* are potentially viable biomarkers [65, 66].

### 2.2 Generalizing Biomarkers

Next we investigated whether our candidate biomarkers could generalize beyond the cohort in which they were discovered. We used the genes found at a given threshold in one cohort, the *source* cohort, to cluster patients in a different cohort, the *target* cohort. Using K-means clustering we partitioned the target population and plotted Kaplan-Meier curves for their progression free survival (PFS) times. The results of this analysis are shown in Figure 3.

**Figure 3:**
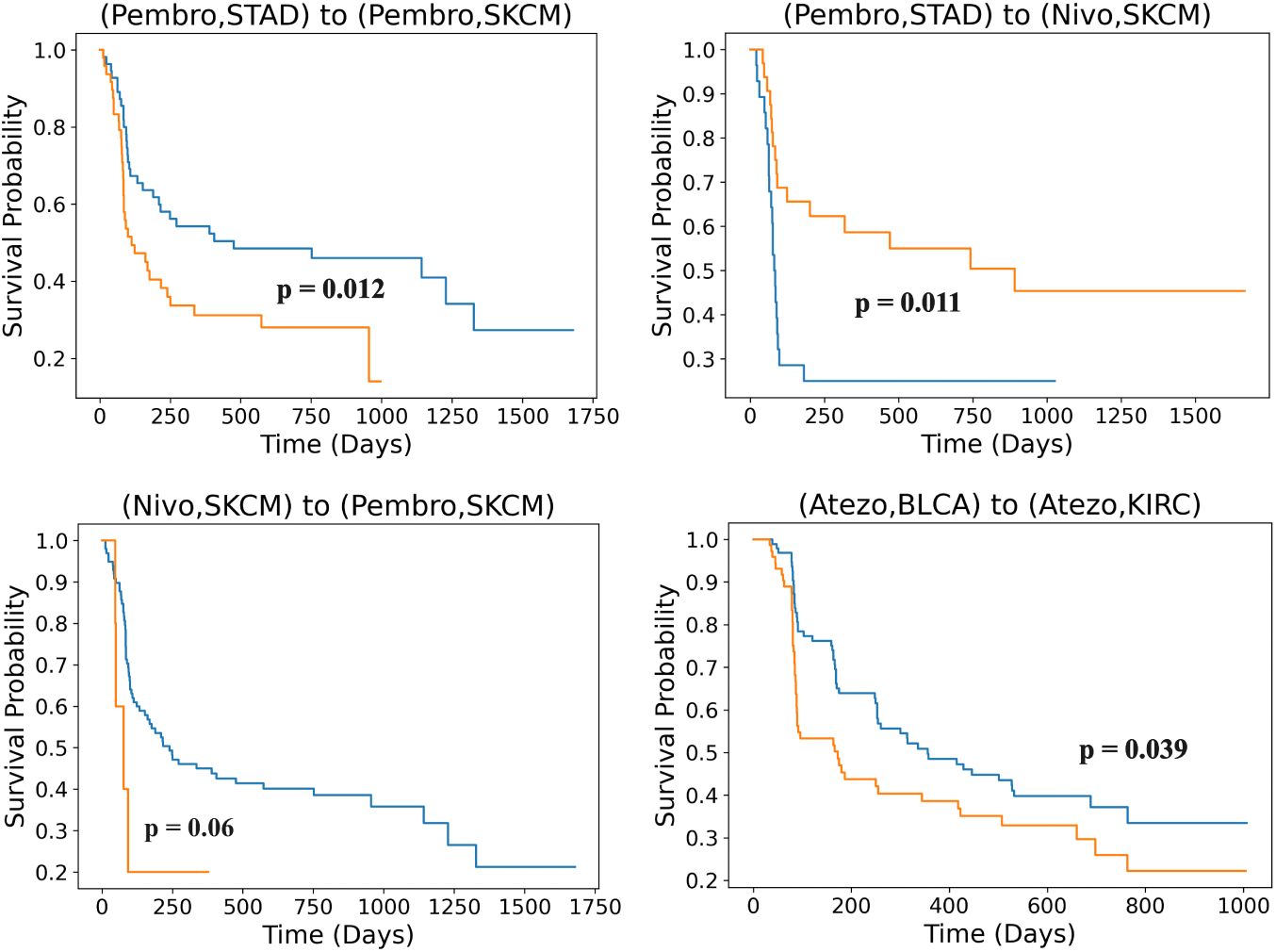
Investigating the ability of biomarkers from one cohort to be used to effectively cluster another cohort. In each case the biomarkers from the source cohort led to clusters which had meaningfully distinct survival curves. The transfer was successful when we analyzed the same drug applied to different tissues (top left, bottom right) between drugs with the same target in the same tissue (bottom left) and drugs with the same target across different tissues (top right). In each case we had a log-rank p-value below 0.1, with most values below 0.05.

We applied this procedure to several transfer scenarios. We investigated the ability of our candidate biomarkers to transfer between cohorts treated with the same drug but with different tissues of origin. We also investigated transfer between drugs but within the same tissue of origin, and across different drugs and different tissues. Transfer between drugs was always performed on drugs with the same target. We used the genes from Figure 2 as candidate biomarkers from the source tissue. Namely, for the (Pembro, STAD) to (Pembro, SKCM) and (Pembro, STAD) to (Nivo, SKCM) transfers we used the genes with edge curvature p-values below 0.05 among the STAD patients treated with pembro which also had Mann-Whitney p-values below 0.1 when comparing the expression distributions between responders and non-responders. In all cases we found two clusters which had survival curves with log-rank p-values below 0.1, with most values below 0.05.

### 2.3 Collating Biomarkers with Rank Aggregation

Since the candidate biomarkers were able to generalize across cancer tissues we endeavored to find aggregated lists of candidate biomarkers for each drug. We expanded our gene sets by looking at edge curvature as well as two ‘scalar’ (vertex-wise) curvatures (for details see Section 4). For each cohort treated by the same drug, *d*, we looked at each tissue *t* for which our pipeline produced candidate biomarkers. For each drug and tissue we created a set *L_dt_* of genes found to have different curvatures (of all kinds) between responders and non-responders. Taking the union of these sets provided a drug-specific gene set *L_d_*.

Then using three feature selection criteria — mutual information [67, 68], the F statistic [69], and the *χ*-squared statistic [70] — we ordered those selected genes by how informative they were in predicting response according to each criterion. For a collection of *T* tissues, this procedure created a total of 3*T* ranked lists which we then harmonized using the Borda count rank aggregation method [71]. Our ranked list of potential biomarkers for each drug are given in Table 2.

**Table 2:**
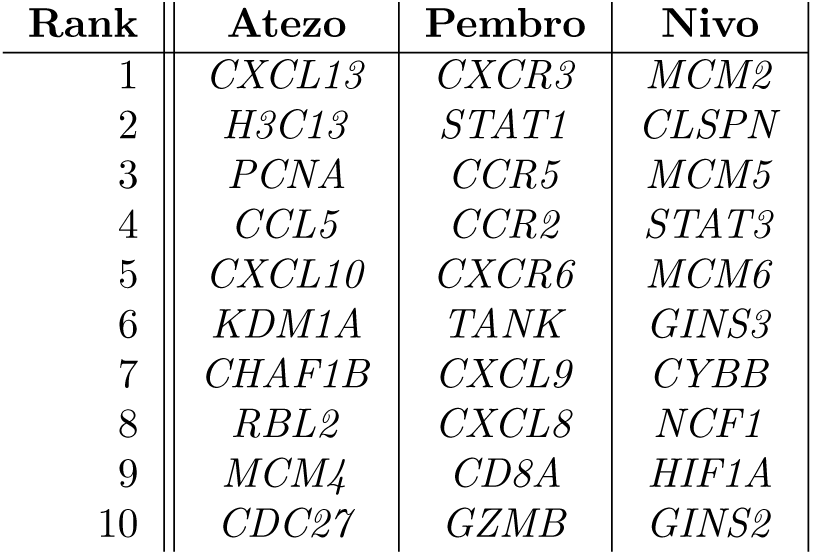
Top 10 drug-specific candidate biomarkers order by aggregated rank. Using edge and node-wise curvature values to create a gene set we then used the Borda count method to aggregate genes into ranked lists.

## 3 Discussion

In this paper we have presented a novel pipeline for discovering biomarkers for response to treatment with immune checkpoint inhibitors. We applied the pipeline to a number of patient cohorts which covered several drugs and tissues. Through rigorous transfer learning experiments we showed that these biomarkers can generalize beyond the setting in which they were found. We harmonized our discovered biomarkers into ranked lists for each drug using the Borda count rank aggregation method.

Our pipeline recovered known biomarkers, many of which have been hypothesized (with experimental support) in publications from the last four years. As such the present study presents supporting *in silico* evidence for those biomarkers, and those biomarkers validate the use of curvature for biomarker discovery. We believe that the genes yielded by our pipeline merit further study in larger data sets and *in vitro* in model organisms or patient derived xenografts [72].

We believe that a natural step to build off the present work would be to develop machine learning models for either response classification or survival forecasting [73, 74]. These experiments require careful construction of sufficiently large datasets corrected for batch effects. The Orcestra database is one such resource [75], but even then the data set sizes may be too small for most machine learning methods. Given a large enough dataset of sufficient quality our candidate biomarkers could be further refined by selecting those which provide the most prognostic capability.

The use of curvature to analyze specific patients could be applied to other data types. For example single cell RNA-seq could be used to compute multiple curvature values per patient [76]. These distributions could then be compared to stratify patient populations. Alternatively, single-sample network inference methods such as Lioness [77] could be used as an alternate method for creating patient-specific networks. Our bootstrapping procedure could be of independent utility in finding differentially expressed genes.

Rank aggregation is worthy of further exploration for analyzing determinants of ICI response. Since there are many types of ICI response biomarkers in the literature that do not involve gene expression, such as tumor mutational burden [11], rank aggregation could be applied to order these different biomarker classes. A recent study performed a similar procedure on data for Crohn’s disease [78]. More sophisticated rank aggregation methods, such as HodgeRank, would likely need to be adopted in exploration of multi-omic ICI response biomarkers [79].

We close by observing that our transfer clustering was not effective when performed with scalar curvatures. This disparity suggests that it is *pairwise* interactions that are important. We believe this supports the use of PPI information in biological inquiry and points to the potential for higher-order interactions to serve as biomarkers [80, 81].

## 4 Methods

### Data Collection & Preprocessing

Our protein-protein interaction (PPI) network was constructed from the collection of interactions in the STRING database [82, 83]. We downloaded the 12th version of the database from the STRING website https://string-db.org. STRING stores interactions as triples of the form (protein 1, protein 2, combined score) where the proteins are named with Ensembl protein IDs [84, 85] and the combined score is a numerical measure of confidence in the given interaction. The combined score takes values from 0 to 1000 with 1000 indicating the greatest possible confidence. We filtered the interactions to only those between proteins with Ensembl IDs which could be uniquely mapped to a HUGO symbol [86] and with combined scores greater than or equal to 950. The resulting graph, *G_♮_* consisted of vertices corresponding to each of the remaining proteins with undirected, unweighted edges corresponding to the remaining interactions.

We downloaded all the ICI data from the CRI-iAtlas (CRI) [87]. We selected only the patient samples collected before any ICI was administered and chose to ignore glioblastoma patients to avoid complications due to the blood-brain barrier [88]. In these selected samples we kept only those genes which had expression measured in both transcripts per million (TPM) as well as raw read counts. We further filtered the list of genes to those which could be mapped to a node in *G_♮_*. The resulting list contained 19,038 genes.

### Seed Gene Discovery

The first step in our pipeline involved the selection of seed genes for each patient cohort. We found these seed genes with a novel bootstrapping procedure.

For 200 iterations we sampled 80% of the cohort without replacement and used the DESeq2 [89] software with default settings to find genes that were differentially expressed (DE) between responders and non-responders. For each gene *g* and for a collection of significance thresholds *ζ ∈* {0.001, 0.002, 0.01, 0.02, 0.05, 0.1} we kept a count *H_gζ_* of the number of times *g* was considered DE between responders and non-responders with an adjusted p-value less than or equal to *ζ*. At the end of this process we had counters *H_gζ_*

For each fixed *ζ* we used an empirical Bayes beta-bernoulli model [90, 91] to estimate the probability that a gene *g* was truly DE at level *ζ*. We created initial probability estimate 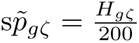 for each gene *g* and used those values to fit the parameters *a*^, 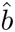 of a beta distribution [92]. The final probability estimates were computed as

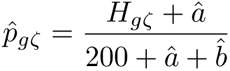

The seed genes were determined by choosing a fixed *ζ* value and keeping those genes *g* with 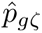 above a threshold *ω*. We excluded KIRC patients treated with nivolumab and SKCM patients treated with pembrolizumab from our analysis as we failed to find any seed genes.

### Constructing Cohort-Specific Graphs

For each cohort consisting of patients with tissue of origin *t* treated with drug *d* we constructed a cohort-specific graph *G_dt_* in three steps. First we manually specified values for *γ* and *ω* and kept the seed genes at *γ* with 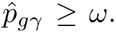 These values were tuned such that all the resulting graphs were of approximately the same order of magnitude (between 100 and 1000 vertices).

As a next step we computed the union of the two-hop neighborhoods (neighbors of neighbors) of each of the seed genes, creating a set *S_dt_*. To complete the construction of *G_dt_* we created the subgraph of *G_♮_* with vertices in *S_dt_* and selecting the largest connected component of this graph. The cohort-specific parameters *ω* and *ζ*, the number of seed genes, the seed gene HUGO symbols, and the graph sizes are given in Appendix A.

### Graph Curvature

Let *G* = (*V, E, w*) be a weighted undirected graph where *V* is a finite set of vertices, *E ⊂ V × V* is a finite set of undirected edges, written {*x, y*}, and *w* is a weight function 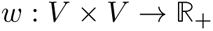 which takes only positive real values. If there is an edge {*x, y*} between a pair of distinct vertices, *x, y*, then we say *x* is *adjacent* to *y*, and write this as *x ∼ y*. This relationship is symmetric, meaning *x ∼ y* implies *y ∼ x*. An edge {*x, y*} is said to be *incident* to its endpoints, *x, y*. For any two vertices *x, y ∈ V* we denote the value of the weight function *w* evaluated at those vertices, as *w*(*x, y*). For any vertex *x*, its degree, *d_x_*, is the sum of the weights of the edges connected to it. That is:

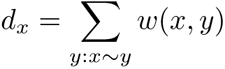

where the sum is taken over all vertices *y* that are adjacent to *x*.

In order to define a probability distribution on each vertex, we first set an *idleness* parameter *α ∈* [0, 1] and assign a probability distribution 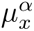 to each vertex *x*:

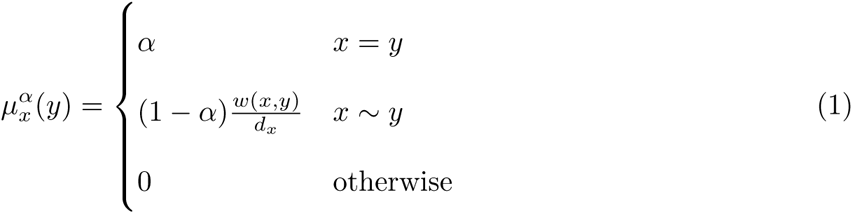

Finally, for an edge *x ∼ y* we define the Ollivier Ricci Curvature (ORC) [17, 93] with idleness *α*, written *κ^α^*(*x, y*) as:

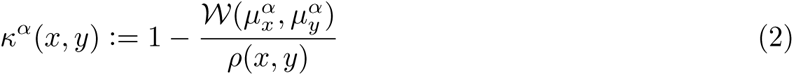

where *ρ* is the length of the shortest path between *x* and *y* and *W* is the Wasserstein distance [94].

As the ORC is defined on *edges*, one can easily define a *scalar* (vertex-wise) *curvature* by summing over the edges incident to a vertex [25]. Specifically, the *scalar* curvature of a vertex *x*, for idleness parameter *α* is given by:

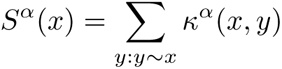

We also consider a *normalized* scalar curvature

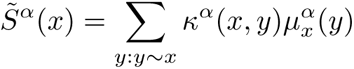

### Transfer Clustering

For our transfer learning experiments we selected the genes from the source cohort and restricted the expression measurements (in TPM) in the target cohort to those genes. We then log-transformed the measurements (log_2_(*TPM* + 1)) and centered and scaled the data so each gene had zero mean and unit variance. To cluster the patients we performed *k*-means [73] clustering for *k ∈* {2, 3, 4}. We chose the *k* for which the silhouette score [95] was largest.

### Assigning Patient Specific Weights

For a given sample *i* let ***x***^(^*^i^*^)^ be the vector of gene expression for that sample where we denote by 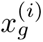 the expression value (in TPM) of gene *g* in sample *i*. We use ***x***^(^*^i^*^)^ to assign weights to the underlying cohort-specific graph via the so-called “principle of mass action” [96]. Concretely, the weight *w*^(^*^i^*^)^(*r, s*) between genes *r* and *s* is given by:

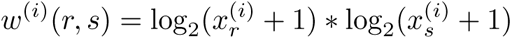

### Rank Aggregation

Rank aggregation is, as the name suggests, a collection of techniques for merging orderings of the same set of objects to one common list. An intuitive example is ordering a collection of people’s individual preferences on a restaurant menu into a single ordering of the menu items from “best overall” to “worst overall.”

To apply these techniques we need to create a collection of ordered lists of genes for each drug *d*. To do this we first curated a set of candidate genes *U_d_* as the union of genes with edge, scalar, and normalized scalar curvature distributions that had Mann-Whitney *p*-values, adjusted for multiple hypothesis testing, below a certain threshold (see Algorithm 1) for each tissue of origin treated with *d*.

Using the python software scikit-learn [97], specifically the function SelectKBest, we ordered *U_d_* from best to worst in terms of three criteria: mutual information [67, 68], the F-test statistic based on analysis of variance [69], and the *χ*-squared statistic [70]. Each of these measures looks at the genes in *U_d_* individually and computes a measure of statistical dependence between expression levels of that gene and patient response. For each of these measures and for each tissue for which we had data (including those for which our pipeline did not find seed genes) we computed orderings for each of these measures. Thus for a total of *T* tissues we had 3*T* ordered lists.

To put these lists together we use a computationally efficient method known as the Borda count. Suppose *U_d_* has *M* genes. For each list *L* gene *g* is assigned a *positional score* based on where it is in the list. If gene *g* is at position *i* in list *L* the score of gene *g* in list *L* is *s*(*g, L*) = *M − i*. We average the scores across all the lists to get a score value for each gene:

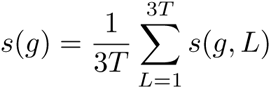

and sorted the genes in decreasing order of *s*(*g*).

### Implementation Details

All code for the experiments in this paper was written in python and R and is available at github.com/jbannon/ORCBiomarkers.

## Acknowledgements

The authors are grateful to the support from NYU during the research that lead to this document. We are also thankful to Benjamin Haibe-Kains for many useful discussions.

## Data Availability Statement

The data needed to recreate the experiments in this paper is publicly available at the CRI i-Atlas portal https://cri-iatlas.org and the STRING database https://string-db.org. The curvature values and p-values are freely available to download at https://zenodo.org/records/13377138.

## Author Contributions

JB, CC, and BM conceived of the experiments. JB wrote the code and performed the analysis and drafted the manuscript. JB, CC, and BM edited the manuscript. All authors have read the manuscript and approve of its submission.

## A Supplementary Material

### A.1 Method Details

Here we provide an algorithmic presentation of the manner with which we found curvature-derived biomarkers using both vertex and edge curvatures.

#### Algorithm 1 Curvature-Based Gene Discovery

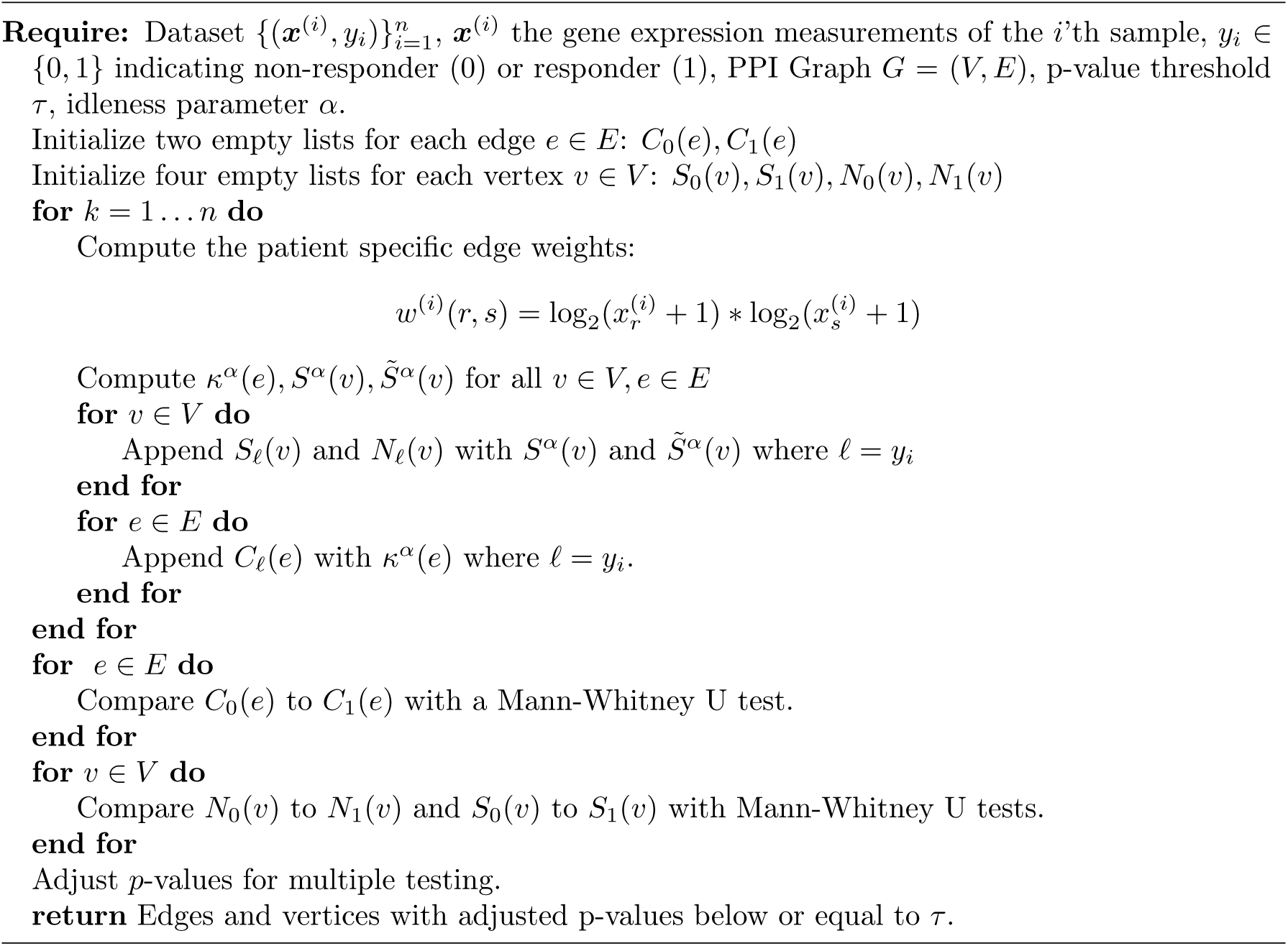

### A.2 Cohort-Specific Graph Construction

This subsection provides information on the sizes of the cohort-specific graphs and how they were constructed. We first give the parameters *ζ* and *ω* for each cohort:

- **Atezo - BLCA:** *ζ* = 0.002*, ω* = 0.9
- **Atezo - KIRC:** *ζ* = 0.05*, ω* = 0.9
- **Nivo - SKCM:** *ζ* = 0.02*, ω* = 0.9
- **Pembro - STAD:** *ζ* = 0.002*, ω* = 0.9

The resulting graph sizes are given in Table 3 and the seed genes are listed below.

**Table 3:** The size of each cohort-specific graph used in our analysis.

| Drug | Tissue | Graph Size | Num. Seed Genes | Threshold $\zeta$ |
| --- | --- | --- | --- | --- |
| Atezo | BLCA | $\vert V\vert = 262, \vert E\vert = 1,263$ | 38 | 0.002 |
| Atezo | KIRC | $\vert V\vert = 589, \vert E\vert = 3,025$ | 44 | 0.05 |
| Pembro | STAD | $\vert V\vert = 166, \vert E\vert = 868$ | 4 | 0.002 |
| Nivo | SKCM | $\vert V\vert = 784, \vert E\vert = 4,414$ | 42 | 0.02 |

**Atezo - BLCA:** MYBL2, REXO5, SLC3A1, CYP4A11, MMP7, ACSM2B, SLC12A1, KRT1, FGA, ALB, SLITRK5, TENM2, PRSS2, HP, SFTPB, ALDOB, PTHLH, HABP2, KNG1, ORM1, HRG, APOA1, C4BPA, PHF6, DPYSL5, BNC1, SLC34A2, C8B, VNN1,MMP10, ACSM2A, FGB, IL1RL1, FGL1, FXYD2, MYOM3, PLG, SERPINA5.

**Atezo - KIRC:** CSRP3, KLRC1, MYPN, MUC5B, SERPINB4,BPIFB1, NRAP, HBG2, EGR3, MYLPF, ZAN, XIRP2, MYH1, UGT2B15, MYBPC1, ACTA1 AKR1B15, MMP1, HAND2, TNNC2, HP, MB, HAO1, PTGS2, MYH8, FLG2, CRP, IL1B, MYH2, TRDN, KRT5, MYH7, MYL1, GUCY2C, DSG1, MMP10, SBSN, FGL1, ARG1, KRT23, MYL2, UGT1A10, CKM, DSC3.

**Pembro - STAD:** WARS1, CXCL10, SYCE2, CXCL11.

**Nivo - SKCM:** TF, PIK3CD, LNP1, CDH10,GAD2, CRISPLD1, SGCD, CLDN7, CDK11B, RBIS, EDNRB, ULK2, RPAIN, PASK, ELF3, STUM, CNTNAP5, LMO3, DDX11, ABCC11, CR-TAC1, MAP6, ZNF781, ELFN2, ZNF727, NANOS1, PLK3, PFN2, TFAP2B, RMDN1, C1ORF116, FAXDC2, ABCA7, USP22, EFHD1, GSTA4, GRIN2B, ZNF704, NME5, GRK2, YWHAE, XIRP1.

### A.3 Additional Results

We lastly present the number of genes from edge, scalar, and normalized scalar curvature that our pipeline discovers at each p-value threshold *τ* . The edge curvatures are given in Table 4, scalar curvatures in Table 5, and normalized scalar cuvatures in Table 6.

**Table 4:** Incremental number of genes yielded as candidate biomarkers from **edge** curvatures. Each column gives the number of *additional* candidate biomarkers discovered by setting *τ* to the given value.

| Drug | Tissue | $\tau = 0.005$ | $\tau = 0.01$ | $\tau = 0.05$ | $\tau = 0.1$ |
| --- | --- | --- | --- | --- | --- |
| **Atezo** | BLCA | 23 | 8 | 50 | 10 |
|  | KIRC | 5 | 1 | 31 | 22 |
| **Nivo** | SKCM | 0 | 0 | 10 | 93 |
| **Pembro** | STAD | 0 | 16 | 12 | 4 |

**Table 5:** Incremental number of genes yielded as candidate biomarkers from **scalar** curvatures. Each column gives the number of *additional* candidate biomarkers discovered by setting *τ* to the given value.

| Drug | Tissue | $\tau = 0.005$ | $\tau = 0.01$ | $\tau = 0.05$ | $\tau = 0.1$ |
| --- | --- | --- | --- | --- | --- |
| **Atezo** | BLCA | 1 | 6 | 39 | 14 |
|  | KIRC | 1 | 0 | 24 | 22 |
| **Nivo** | SKCM | 0 | 2 | 0 | 28 |
| **Pembro** | STAD | 0 | 0 | 3 | 5 |

**Table 6:** Incremental number of genes yielded as candidate biomarkers from **normalized scalar** curvatures. Each column gives the number of *additional* candidate biomarkers discovered by setting *τ* to the given value.

| Drug | Tissue | $\tau = 0.005$ | $\tau = 0.01$ | $\tau = 0.05$ | $\tau = 0.1$ |
| --- | --- | --- | --- | --- | --- |
| **Atezo** | BLCA | 4 | 3 | 27 | 18 |
|  | KIRC | 1 | 1 | 25 | 10 |
| **Nivo** | SKCM | 0 | 2 | 0 | 49 |
| **Pembro** | STAD | 0 | 0 | 4 | 9 |

## Notes

https://zenodo.org/records/13377138

### Competing Interest Statement

The authors have declared no competing interest.

